# Perplexity about periodicity repeats perpetually: A response to Brookshire

**DOI:** 10.1101/2022.09.26.509017

**Authors:** Daniele Re, Tommaso Tosato, Pascal Fries, Ayelet N. Landau

## Abstract

Brookshire (2022) claims that previous analyses of periodicity in detection performance after a reset event suffer from extreme false-positive rates. Here we show that this conclusion is based on an incorrect implemention of a null-hypothesis of aperiodicity, and that a correct implementation confirms low false-positive rates. Furthermore, we clarify that the previously used method of shuffling-in-time, and thereby shuffling-in-phase, cleanly implements the null hypothesis of no temporal structure after the reset, and thereby of no phase locking to the reset. Moving from a corresponding phase-locking spectrum to an inference on the periodicity of the underlying process can be accomplished by parameterizing the spectrum. This can separate periodic from non-periodic components, and quantify the strength of periodicity.

## Introduction

Brookshire (2022) revisited reports of rhythmicity in detection performance (e.g., Landau and Fries, 2012; Fiebelkorn et al., 2013), and concluded that formerly employed methods lead to excessive false-positive rates. Previous studies had presented, per trial, one reset event (a flash), followed by one randomly timed probe, and had recorded the behavioral response (hit or miss). Across many trials, the reset-aligned accuracy time course (ATC) was calculated. The ATC was then Fourier transformed, and the resulting spectrum compared to spectra obtained after randomly pairing, across trials, behavioral reports and probe time points, i.e., after “shuffling-in-time”. This procedure tests for temporal structure. Brookshire makes the valuable point that rejecting the null hypothesis of no temporal structure does not unequivocally demonstrate the presence of periodic structure, and therefore argues that the null hypothesis should consist of a temporal structure that is aperiodic.

### The calculation of false positives - a single noisy time course is not noisy enough

Brookshire’s implementation of the aperiodic null hypothesis is based on different types of noise processes, primarily the first-order autoregressive (AR(1)) process and its special case, the random walk. In an AR(1) process, the signal at time t is the sum of a specified fraction of the signal at time t-1 plus a random step (Figure 1A). When many realizations of an AR(1) process are Fourier transformed, their average spectrum decays monotonically with frequency according to 1/f^n^, without peaks indicative of periodicity (Figure 1B, left). However, single realizations of an AR(1) process often yield spectra that do not decline monotonically with frequency and thus have spectral peaks (Figure 1B, right). Despite this fact, Brookshire simulates the ATC on the basis of a single AR(1) realization; this realization is taken as a probability time course, and probabilistic draws from it generate the hits and misses of all trials (and *all subjects*; Figure 1C, D); the resulting ATC is then analyzed with the shuffling-in-time statistics, often yielding significant results for some frequency bins (Figure 1E-G). Brookshire argues that these results should be considered false positives, because the AR(1) process is aperiodic. However, as explained above, this does not hold for single AR(1) realizations. When we use the code provided with Brookshire (2022) and modify it to implement separate AR(1) realizations for each trial of each subject (Figure 2A), or even just for each subject (Figure 2B), false positives are substantially diminished.

**Figure 1:**
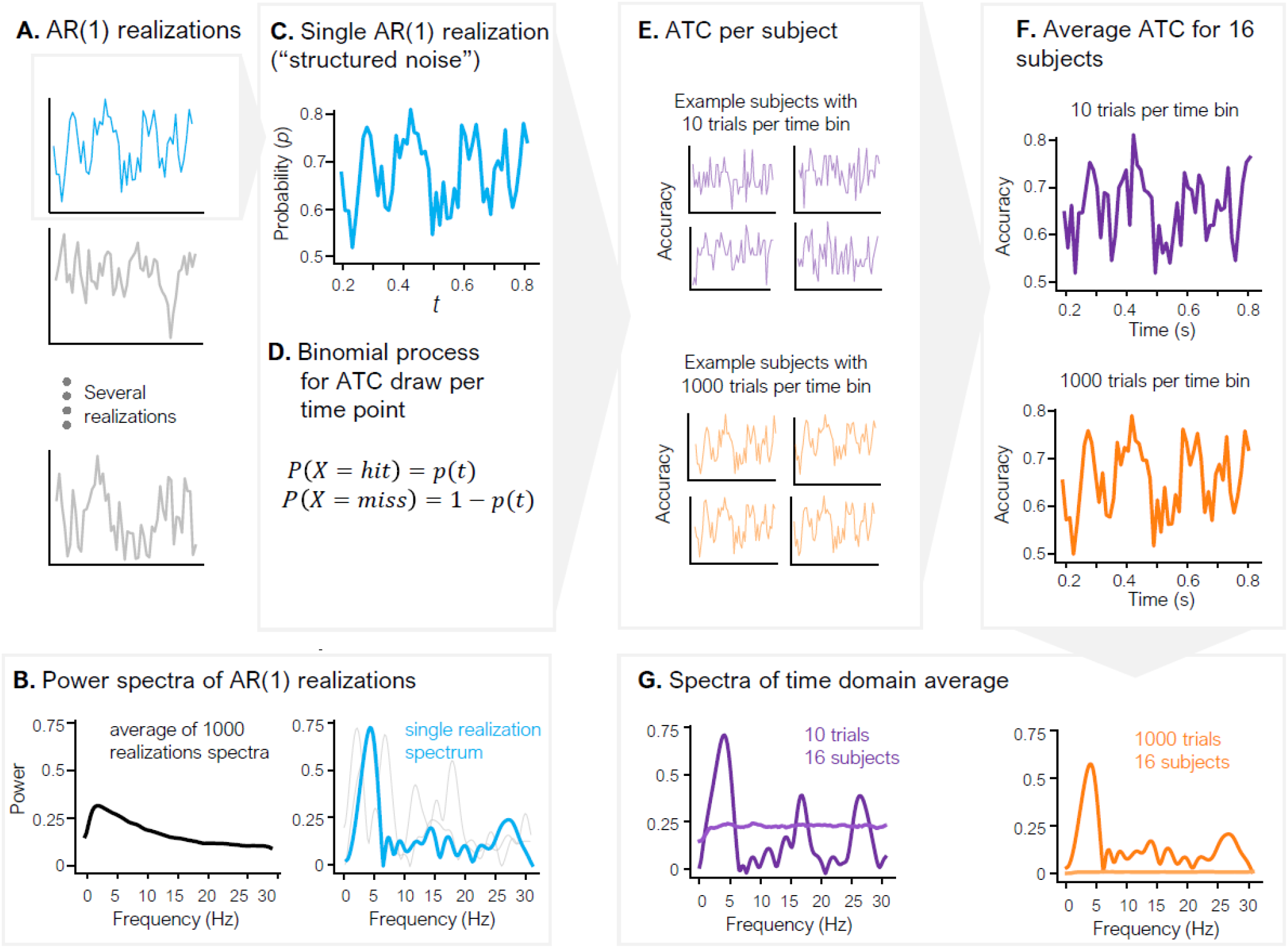
Illustration of the null-hypothesis implementation proposed by Brookshire (2022). (A) Several realizations of an AR(1) process. (B) Left: Power spectrum averaged over 1000 realizations. This average spectrum declines monotonically, except at its low-frequency end where it shows the effect of linear detrending. Right: Power spectra of single realizations, showing clear peaks. (C) The single realization used as a probability function for hit and misses underlying the ATC. Brookshire refers to this probability function as “structured noise”. (D) A binomial process is used to draw the single-trial outcome from the probability distribution in C for each time bin *t*. (E) The outcomes are averaged over trials to obtain the ATC for each subject. (F) The ATCs averaged over subjects are shown for different numbers of trials per time point (10 and 1000, respectively). Average ATCs are similar to the single AR(1) realization shown in C, more so, the more trials are included. (G) ATC power spectra, and corresponding significance thresholds (dashed), for 10 (purple) and 1000 (orange) trials per time bin. Increasing trial numbers lead to increasing significance, contrary to what is expected from a noise process.

**Figure 2:**
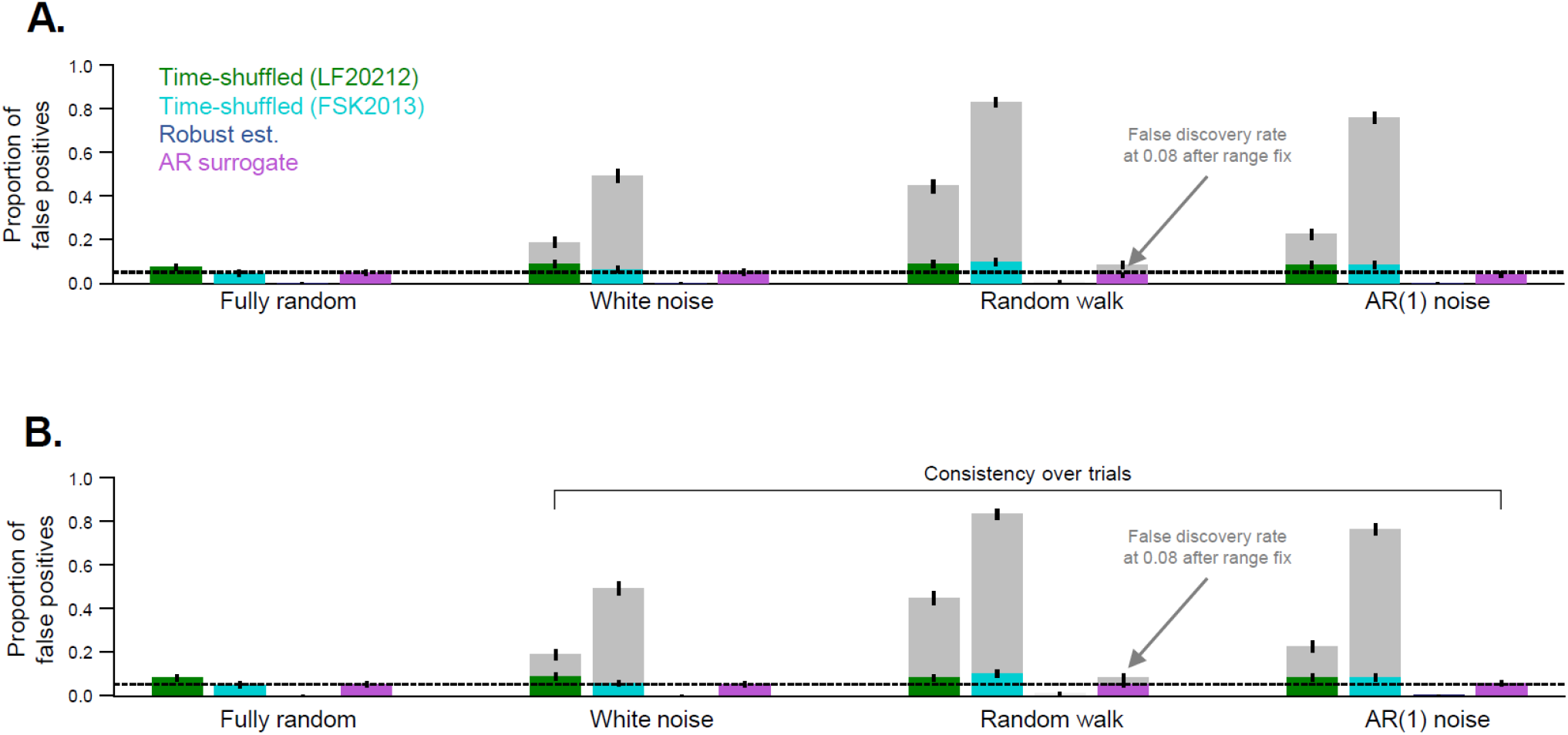
Replotting Figure 3a from Brookshire (2022) using provided analysis code. The gray bars show false-positive rates reported in Brookshire (2022), where trials and subjects were drawn from a single realization of the chosen noise process (except for the Fully random condition). The colored bars are based on the same analysis, only implementing separate realizations per trial (A) or per subject (B), which resulted in false-positive rates close to 0.05, with negligible differences between methods. Although separate realizations should be used per trial (panel A), even the use of separate realizations merely at the level of each subject (panel B) is sufficient to have low false-positive rates in all analysis methods. Note that the gray bars for the AR surrogate include a normalization step, which leads to higher false-positive rates, and which was missing in Brookshire’s implementation (see section on “Methods proposed by Brookshire – and their problems”).

The use of a single time course to generate many simulated trials (and many subjects) trivially leads to phase-locked modulation of simulated behavior. The hits and misses generated in single trials are just (very) noisy replications of the single time course. If this time course is not entirely flat, then it has some temporal structure, and the noisy replications of this temporal structure across trials are equivalent to phase locking of the trials to the reset event. Thus, phase-locking metrics as used in Landau and Fries (2012) should and do actually provide significant results in this case. The significance increases when more trials are simulated (Figure 1G), demonstrating that many draws of a single time course are not an implementation of “structured noise” as claimed by Brookshire.

Note that several previous studies modeled e.g. evidence accumulation as AR(1) process (drift diffusion), but they consistently implemented separate AR(1) realizations per trial (Ratcliff and McKoon, 2008; Shadlen and Kiani, 2013)). Other studies did use one function to model trends in trial-averaged behavioral time courses, but they used deterministic processes, such as Gaussian or exponential functions (Grabenhorst et al., 2019; Grabenhorst et al., 2021), and not stochastic ones, such as AR(1) or random walk.

### From spectra to interpretation

It is important to clarify what shuffling-in-time actually tests. Shuffling-in-time followed by Fourier transformation is equivalent to shuffling-in-phase. If statistical tests based on shuffling-in-phase are significant for a given frequency bin, this means that there is significant phase locking (to the reset event) at that frequency bin. An isolated significant frequency bin in a phase-locking spectrum is consistent with a periodicity in a frequency band including this frequency, i.e. with a spectrum containing a distinct peak. Yet, it is also consistent with a different spectral pattern that is not indicative of periodicity. To move from a phase-locking spectrum to the inference on a likely underlying, periodic or non-periodic, process, one needs to consider the shape of the entire spectrum or at least of a substantial part of the spectrum (Tosato et al., 2022). This interpretation of the spectrum can be achieved by, e.g., parameterizing the spectrum (Donoghue et al., 2020). Such parameterization can objectively separate periodic from non-periodic components, and quantify the strength of the observed periodicity.

When periodicity has been established, the evidence can be further strengthened by replication, e.g. across different conditions within one study (for a similar approach, see Vinck et al., 2022). Indeed, several studies have found that different experimental conditions produce phase locking to the reset event at very similar frequencies (Landau and Fries, 2012; Zhang et al., 2019).

### Methods proposed by Brookshire – and their problems

Brookshire (2022) proposes two methods for analyzing behavioral time courses, namely “AR surrogate” and “robust estimation”, which are presented as having better detection ratio (the ratio of true positives to false positives). The AR surrogate method models the empirical ATC with an AR(1) process, and then uses this AR(1) process to generate surrogate ATCs, which form the basis for statistical testing. This method does generate multiple realizations of the AR(1) process. However, the surrogate ATCs are scaled using the standard deviation of the noise, which unfortunately causes their values to exceed the range of possible detection rates, i.e., 0 to 1. This leads to an inflation of the power of the surrogate data compared to realistically simulated and empirical data. As a result, Brookshire (2022) reports a very low false-positive rate with this method, which leads to falsely high detection ratios. When the scaling is corrected, false-positive rate is higher (Figure 2, arrow; 0.08 instead of 0.03 in Brookshire (2022)), and detection ratio is slightly lower.

The second method proposed, the robust estimation method, is presented as having an acceptable detection ratio. However, the true-positive rate of this methodology is unacceptably low (Figure 3; as pointed out by several commentaries on this work, e.g., Fiebelkorn, 2022; Vinck et al., 2022). This fact is masked in the detection ratios by false-positive rates approaching zero. Figure 3 shows the false- and the true-positive rate for our method as well as the two methods proposed by Brookshire. We simulated a periodic modulation with a frequency of 4 Hz and with modulation depths (defined as in Brookshire (2022)) of 0.3, 0.2, and 0.1, similar to empirically observed modulation depths (Busch et al., 2009; Landau et al., 2015; Benedetto and Morrone, 2017; Tomassini et al., 2017; Re et al., 2019). On these simulated data, all methods were tested, and the methods proposed by Brookshire (2022) suffer from very low true-positive rates (Figure 3).

**Figure 3:**
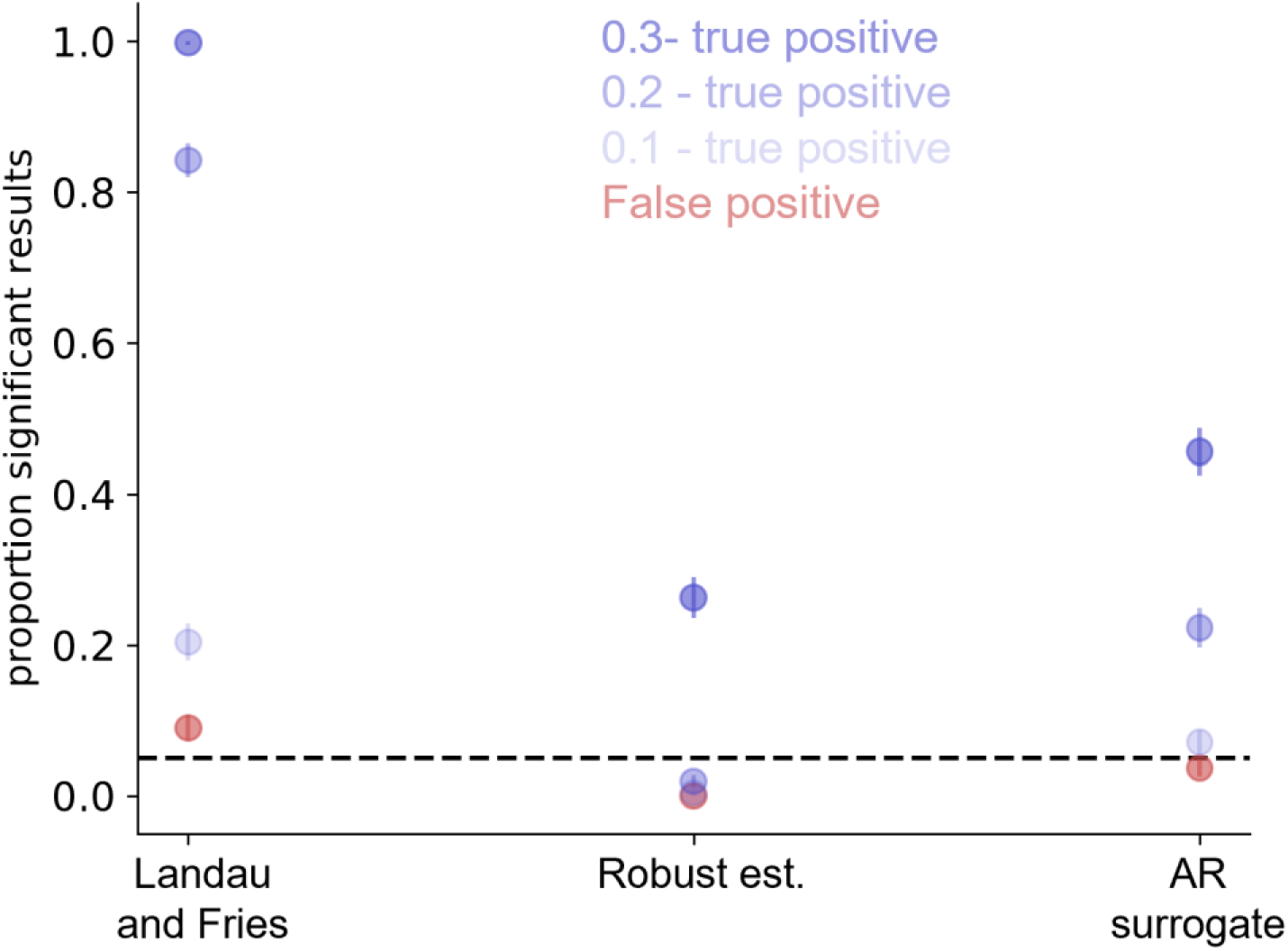
True-positive and false-positive rates for different analysis methods. We simulated a probability time course characterized for all trials by a 4 Hz sinusoidal modulation, and additionally added different random-walk noise, per trial. We simulated 3 conditions with sinusoidal modulation depths (defined as in Brookshire (2022)) of 0.3, 0.2 or 0.1, and one condition without modulation. For the conditions with modulation, the y-axis reflects the true-positive rate, and for the condition without modulation the false-positive rate. As in figure 2, a previously used method (Landau and Fries, 2012) results in low false-positive rates and reasonable true-positive rates. Robust estimation and AR surrogate on the other hand show a true-positive rate below 0.5 for all conditions.

## Conclusion

Brookshire’s main claim of extreme false-positive rates in previous analyses is unfounded. Previous analyses correctly tested for phase locking per frequency. Moving from a phase-locking spectrum to an inference on (the periodicity of) the underlying process can proceed by parameterizing the phase-locking spectrum - a fruitful endeavor for future work.

## Author contributions statement

**Daniele Re:** Conceptualization, Methodology, Software, Formal analysis, Visualization, Writing – Original Draft, Writing – Review & Editing. **Tommaso Tosato:** Conceptualization, Methodology, Software, Formal analysis, Visualization, Writing – Original Draft, Writing – Review & Editing. **Ayelet Landau:** Conceptualization, Methodology, Formal analysis, Visualization, Writing – Original Draft, Writing – Review & Editing, Supervision, Project administration, Funding acquisition. **Pascal Fries:** Conceptualization, Methodology, Formal analysis, Writing – Original Draft, Writing – Review & Editing, Supervision, Project administration, Funding acquisition.

